# A topological perspective on the dual nature of the neural state space and the correlation structure

**DOI:** 10.1101/2023.10.17.562775

**Authors:** Melvin Vaupel, Erik Hermansen, Benjamin A. Dunn

## Abstract

When analysing neural activity, one often studies either the neural correlations or the state space of population vectors. However, the reason for choosing one over the other is seldom discussed. Here, with methods from the mathematical field of topology, we compare these approaches and propose using both for unsupervised inference of neural representations. If the recorded neurons have convex receptive fields on a single covariate space, there is a duality between the topological signatures derived from correlations on the one hand and population vectors on the other hand. However, in the presence of multiple neural modules with non convex receptive fields, this duality breaks down. We explain how to leverage complementary information derived from both approaches to sucessfully characterize the represented covariate spaces directly from the data also under these challenging circumstances. Furthermore, we prove appropriate reconstruction results and showcase applications to multiple neural datasets from various brain regions and diverse neural modules.

## 1 Introduction

In modern neuroscience, the electrical activity of thousands of neurons can be recorded over extended periods of time (Fig. 1a) [1]. Commonly, external covariates, thought to be relevant for the experiment, are recorded alongside the neural activity and one may study how the spiking behaviour of each neuron is modulated by these covariates. Indeed, many types of neurons exhibit distinct peaked response patterns in relation to specific physical variables, such as spatial location or visual stimuli have been identified [2–4]. On the other hand, it is not always feasible or desirable to study neural activity solely as a response to a recorded stimulus; the relevant covariates may be unavailable or unknown for recording. Assuming a covariate may also overshadow other functions of the network. It is thus necessary to consider methods for studying the neural activity in itself, without a priori assumptions about the external covariates.

**Fig. 1:**
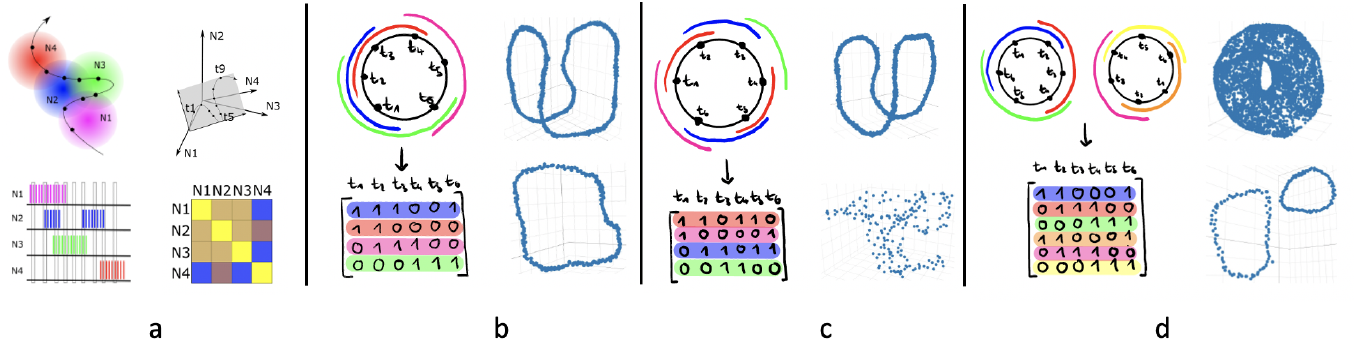
Two approaches of studying neural activity. **a**. Top left: A path through a twodimensional (2-D) covariate space and receptive fields of four neurons, giving rise to spike trains (bottom left). By aligning and binning these, one may construct a space of population vectors (top right) - *column space* - or analyse the neurons’ correlation structure (bottom right) - *row space*. **b.-d**. Simulated neural activity with distinct properties (left) and 3-D UMAP projections [5] (right) of the column space (top) and the row space (bottom). **b**. For neurons with single-peaked receptive fields the circular covariate space is recovered from both the population vectors and the correlations. **c**. For multi-peaked fields the population vector space recovers the circular shape while the correlation structure does not. **d**. For neural activity from two neural populations with single-peaked fields on separate circular covariate spaces we see a torus (the product of two circles) in the population vector space, while the neural correlation space disentangles both representations, showing two circles.

Assuming *N* neurons are recorded during *T* time points, the neural activity can be represented as a matrix *X*, where *X*_*ij*_ represents the activity level of neuron *i* at time *j*. This matrix may be binarized, so that 1 indicates activity and 0 inactivity. The goal is then to study neural function solely based on information we can gather from this neural activity matrix. We will, in the following, take a closer at two groups of existing methodologies: 1) studying the *columns* of the matrix and 2) studying its *rows*.

### Studying columns

The columns of *X* comprise *T* vectors of dimension *N*, each describing the activity state of a population of neurons at a given time point. These are therefore called *population vectors* or *population codes*. We can define a metric on the space of observed population vectors in which two vectors are close if they share many coactive neurons [6]. It is hypothesized that the population vectors, together with this metric, constitute a discrete sample from a continuous underlying space of neural states - the population’s *state space*. This space if often found to be low-dimensional [7] and the constrained activity has been suggested to arise from the connectivity of the networks [8] or through coordinated neural responses to distinct covariates [9]. By studying the shape and geometry of these spaces, e.g., through dimensionality-reduction tools [10], topological data analysis (TDA) [11] or latent variable models [12], we may further our understanding of the function of the population. For instance, in [13] the geometry of the dimension-reduced population activity in the mouse hippocampus reflects both abstract and physical variables during a decision-making task, and the topological structure of the state space of head direction cells and grid cells [2, 14] have been shown to form a 1-D ring and a 2-D toroidal manifold representing head direction and position in space [15–17]. This approach is exemplified in Fig. 2, where we reveal a circular feature in the population activity of orientation-tuned neurons in macaque monkeys [18] (Fig. 2a,b), suggesting a neural representation of the orientations of drifting gratings (Fig. 2d).

**Fig. 2:**
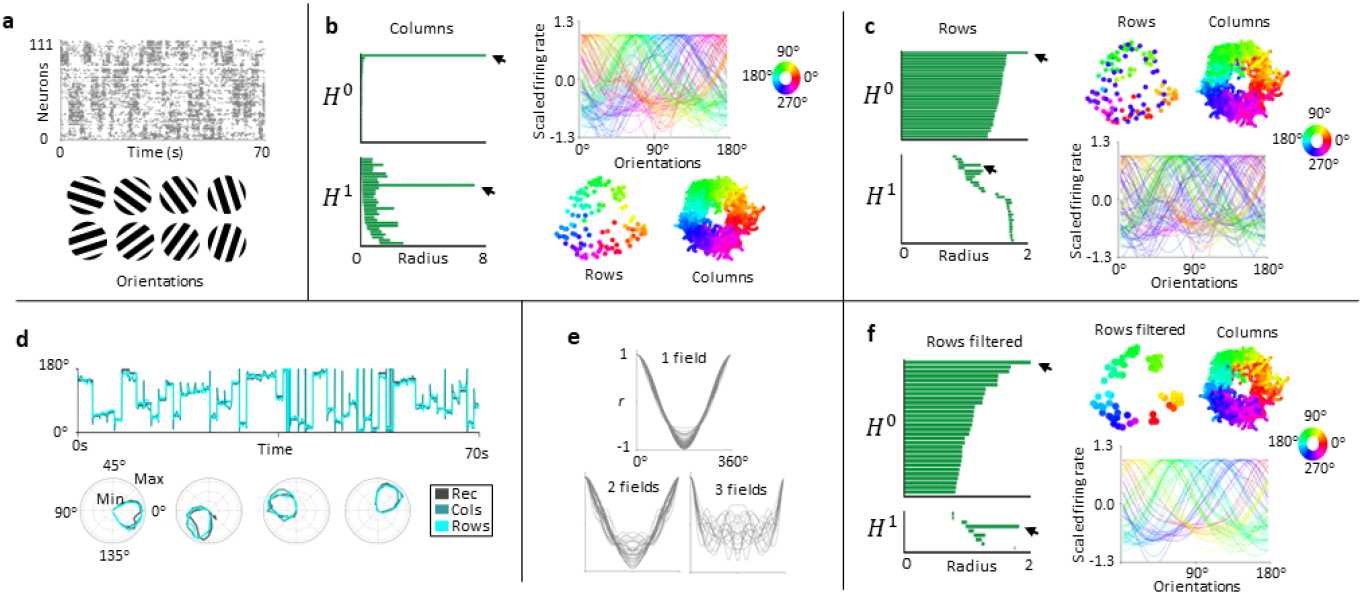
Topological data analysis reveals a ring representation of the activity of orientation-tuned neurons both when studying the rows and the columns of the activity matrix. **a**. Top, spike trains from 111 cells in primary visual cortex in a macaque monkey during presentation of differently oriented, drifting gratings (bottom) [18]. **b**. Left, barcode from applying persistent cohomology to the columns of the activity matrix. The long bars (black arrows) indicate clear expression of a 0- and a 1-D hole, as expected by a circular feature, in the data (*H*^0^ and *H*^1^). Bottom right, UMAP projections of the rows and columns colored according to the neurons’ mean and instantaneous decoded internal representation, respectively. Top right, single cell tuning to grating orientation, colored by mean decoded angle. **c**. Barcodes and decoding as in **b**, studying the rows instead of the columns of the activity matrix. **d**. Tuning of four example neurons (bottom) and snippet of recorded orientations and decoded circular representations (top). **e**. Autocorrelograms of tuning to column-wise decoded neural representation clustered by number of receptive fields observed in each curve. **f**. Barcodes and decoding as in **b**,**c**, analyzing only the cells with single fields as seen in **e**.

### Studying rows

The rows of the matrix *X* give *N* vectors of dimension *T*, often referred to as *spike trains* or *neural codes*, with each vector describing the activity of one neuron across the whole recording. A similarity measure in this space is given by the pairwise correlations of the vectors, in which two neurons are more correlated if there are more time points when they were both active (Fig. 1d). This may be understood as a measure of *functional connectivity* in that high coactivity should indicate a strong computational relation. We can interpret the correlations as weights on a graph, allowing studying the correlation structure on a network level [19–21]. Commonly in large-scale recordings, we find groups of neurons with high inter-correlation. These clusters are often referred to as neural *ensembles* and are presumed to solve specific computational tasks [22–24]. Studying the pairwise correlations of afformentioned orientation-tuned neurons in macaque monkeys, we again observe an underlying circular feature (Fig. 2c-f).

While population vectors and spike trains are extracted as columns and rows of the same data matrix, they may encode complementary information. For instance, different combinations of population vectors can arise from the same correlation structure, and consequently, ‘studying columns’ is more sensitive to the combinatorial possibilities of coactive neurons. On the other hand, the correlation structure incorporates information about how often collections of neurons were co-active, while studies of population vectors are insensitive to such information. Given these differences, the columns and rows are utilised in the analysis of neural data for very different ends. For example, neural ensembles are commonly separated by leveraging the correlation structure, and the population vectors are used to study the computational function of such ensembles.

Despite these differences, the spike trains and population vectors of a neural recording are certainly not independent. It may help to visualize neurons as having receptive fields on the state space of population vectors (Fig. 1). If two receptive fields overlap in more population vectors, the corresponding neurons will have a higher correlation. A formal way of understanding this connection comes from analysing the data matrix *X* through the lens of algebraic topology, a mathematical discipline providing tools to study objects up to continuous deformations. The receptive fields may in certain situations be topologically trivial (contractible), giving rise to a duality between ‘studying columns’ and ‘studying rows’. In other situations this duality can break down and both modes of analysis provide complementary topological information. A major objective of the present paper is to better understand when this happens.

### Outlook

In Section 2, we review why topology is useful for the (unsupervised) inference of neural representations from data. We elaborate why topological tools provide a reasonable framework for such inference and motivate and explain the relevant concepts in a way that is accessible to non-mathematicians. In Section 3 we zoom out and explain two general constructs that associate *weighted simplicial complexes* to neural activity. One of these will represent ‘studying columns’ and the other one ‘studying rows’. We explain how the two underlying simplicial complexes are related through *Dowker duality*, establishing a rigorous link between the two approaches of neural data analysis. Importantly, this duality only holds between the skeleta of the unweighted complexes, while the weights may incorporate additional information. In Section 4 we explain how these constructs may be leveraged for the inference of neural representations and also demonstrate that these methods are useful in the analysis of real neural data. We find that,

1. We may disentangle mixed neural populations using the rows of the data matrix but not with the columns.
2. We may disembiguate non-convex codes using the columns of the data matrix but not with the rows.
3. In recordings with mixed neural populations and non-convex codes we can still infer the correct shapes of the relevant covariate spaces. This requires the combination of information obtained from neural correlation and population vectors.
4. For recordings of single ensembles with convex codes, the duality holds and both weighted simplicial complexes yield the same topological signatures. Conversely we may use such coincidence as a signature for recordings of distinct ensembles with convex codes.

We apply these methods to recordings of the hippocampus, the entorhinal cortex and the primary visual cortex (V1), and determine the underlying topology in population activity of head direction cells, place cells, grid cells and orientation-tuned cells without reference to the respective covariates.

Note, in the appendix, we provide a brief, mathematically more rigorous discussion of the relevant concepts. In particular, we prove two reconstruction results that shed light on how filtrations of Dowker complexes in terms of total weights can be useful for the inference of covariate spaces from activity data of multiple neural modules with non-convex codes.

## 2 Discovering neural representations

In a neural representation, the collective activity of a population of neurons is orchestrated to encode a specific covariate [25]. This means that the properties of the covariate are reflected in activity patterns of the neural activation and can, in principle, be read out by downstream neurons, providing context to subsequent computations. The historically first example of a neural representation was orientation tuning in the visual cortex, where neurons encode the orientation of straight lines in the visual field [3] (Fig. 2a). Each of these neurons displays elevated activity for a small range of orientations, called a *receptive field*, and different neurons respond to different angles. This narrowly peaked response pattern is similarly observed for example in head direction cells [14] and place cells [4]. For the latter, the encoded covariate is the animal’s 2-D spatial environment, and the neurons display 2-D spatial regions of elevated activity often called *place fields*. All of these neural representations were discovered by plotting neural activity against the respective covariate and studying the so obtained *tuning curves*. While this can be an effective tool for studying representations, it comes with limitations. For instance, if the relevant covariate is not known or recordable during the experiment, this approach is naturally not applicable. Thus, an important question of contemporary neuroscience is: what can be derived about the covariate space of a representation directly from the neural recording, without a priori knowledge of the relevant stimulus?

Assume we record the activity of neurons that represent a covariate space *C* with receptive fields as in Fig. 3. We do not know what *C* is a priori, but can derive from the recorded data *X* which receptive fields presumeably overlap, giving a basic outline of *C*. To obtain a more detailed model, we have to make additional assumptions about how the covariate is encoded in the neural activity. For example, we could postulate specific sizes and shapes for the receptive fields or assume that they are uniformly distributed on the covariate space (as is done in [26]).

**Fig. 3:**
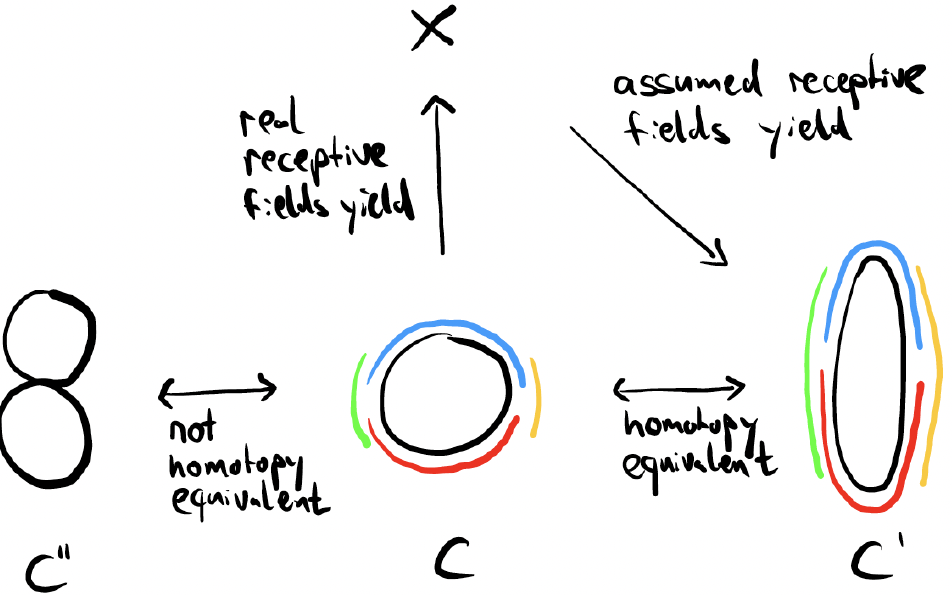
Inferring homotopy type is less sensitive than inferring geometry. The data matrix *X* is a sample of the receptive fields (color-coded) on covariate space *C* (black circle). Assuming a wrong geometric shape of the receptive fields gives a space that is slightly stretched, *C*^*′*^. The space *C*^*′′*^ could not have been inferred from *X*, given the assumed receptive fields.

However, it is often not clear what the right assumptions are. In Fig. 3, the assumption of uniformity would yield the elliptical space *C*^*′*^ instead of the circular *C*. On the other hand, we would not end up with the inferred space *C*^*′′*^ as long as we respect the combinatorics of the overlaps indicated from the recording. In what way is *C*^*′*^ less different from *C* than *C*^*′′*^ is? One possible answer is that we can deform *C* into *C*^*′*^ without ‘gluing or ripping’ parts of the space, while such a deformation does not exist between *C* and *C*^*′′*^. The mathematical field of *topology* makes the notion of deforming spaces without gluing or ripping precise, and two spaces that admit deforming into one another in this way are called *homotopy equivalent*. Thus, if we want to characterize a represented covariate space directly from a neural recording, but have no detailed knowledge of how the covariate is encoded, then we should first try to characterize it with properties that are invariant under homotopy equivalence.

In existing literature, *homology* has often been used as such a homotopy invariant property [16, 27, 28]. Intuitively, the *n*’th homology of a space measures the number of *n*-D holes in it. A sphere has a 2-D hole but no 1-D hole and a torus has a 2-D hole and in addition to two 1-D holes. Indeed, if we can deform two spaces into one another (without gluing or ripping), they will have the same number of *n*-D holes. This tells us, for instance, that the sphere and the torus cannot be homotopy equivalent because they differ in their number of *n*-D holes. In the above example, the space *C* (circle) and *C*^*′*^ (ellipse) both have a single 1-D hole, while *C*^*′′*^ has two 1-D holes, thus the homology of *C*^*′′*^ is different from that of *C* and *C*^*′*^.

### Remark 2.1.

That a space can be recovered up to homotopy equivalence from how receptive fields overlap is due to a result called the *nerve lemma*. This result requires some assumptions on how the receptive fields cover the space, essentially guarenteeing that the combinatorics of how they overlap encode all topologically relevant information (see Appendix A). A sufficient requirement is that all receptive fields are convex subsets of some Euclidean space. However, In Section 4 we will see that this requirement is not always necessary for the topological inference to work and in Appendix A we prove an appropriate reconstruction result.

## 3 Topology and spiking data

The takeaway of Section 2 was that, in characterizing a represented covariate space directly from neural recordings, it is often a good idea to first ask for properties invariant under homotopy equivalence. This can for instance be the homology of the covariate spaces. How do we get from the neural activity data to a space we can calculate the homology of? In the Introduction, we discussed how to approach the analysis of neural data by either considering correlations between neurons or by investigating the space of population vectors. We will now explain two related constructs which allow the inference of homotopy equivalent properties – the *correlation complex* and the *population vector complex*. Comparing them will exhibit a duality between the two modes of analysis.

### Correlation complex

Consider the binarized neural activity matrix, *X*, as shown in Fig. 4. We may represent each column (population vector) as a point and group these into collections based on the neurons they feature as being active. For example, the population vectors *p*_1_, *p*_2_ and *p*_3_ are in one collection because the (green) neuron *n*_2_ is active in each of them. We may then assign a geometric object to this data, called a simplicial complex. This complex has the rows of the matrix *X* (the neurons) as vertices. We connect two vertices by an edge if the corresponding neurons were active in the same population vector – e.g., *n*_1_ and *n*_3_ are connected because both were active in *p*_4_. Larger multiples of neurons are connected by higher simplices (triangles, tetraheda, …). For example *n*_1_, *n*_2_ and *n*_4_ are connected by a 2-simplex (triangle) because they overlap in the population vector *p*_1_. Furthermore, we may assign to each simplex a weight (called the *total weight*, see Appendix A), counting the number of population vectors that the respective neurons share. For example, *n*_2_ and *n*_4_ both appear in *p*_1_, *p*_3_ and *p*_5_ and the corresponding edge has weight 3. Similarly *n*_1_ and *n*_3_ only co-appear in one population vector *p*_4_, so their edge is weighted 1. Because two neurons are more correlated if they are active in many overlapping population vectors, we can think of the weights on the edges as (unnormalized) correlations. Weights on higher simplices denote correlations between more than two neurons. The simplicial complex is thus a geometric representation of a higher-order correlation matrix and we call the simplicial complex thus constructed the *correlation complex*.

**Fig. 4:**
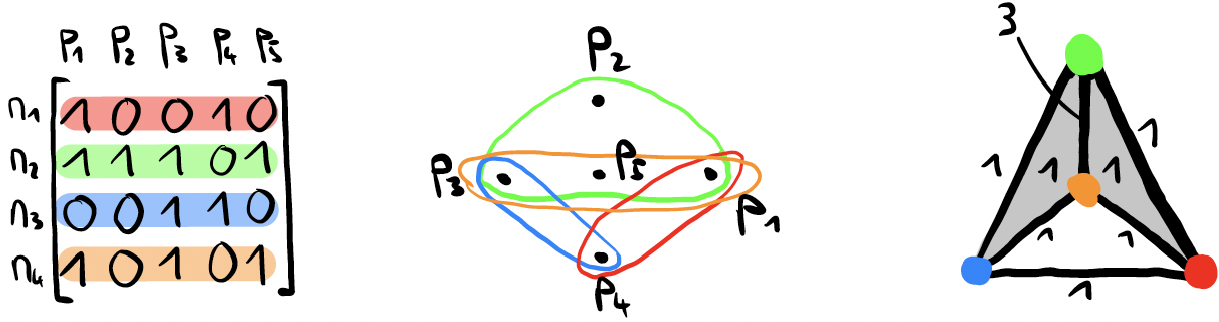
The correlation complex (right) is constructed by grouping the population vectors in the recorded neural data (left) containing the same neuron(s) (middle). Color indicates neuron identity. The weights assigned to each simplex count the number of points that are in the overlap of the respective collections.

### Population vector complex

We now swap the roles of the rows and columns in the above construction, such that every row (i.e., every neuron) defines a point (Fig. 5). The points are then grouped according to the population vectors that feature the corresponding neurons as active. For instance, *n*_1_, *n*_2_ and *n*_4_ are grouped together because they all feature in the red population vector *p*_1_. This gives rise to a simplicial complex where the vertices are the population vectors and the weights on the simplices between them indicate the number of shared neurons. We call this complex the *population vector complex*, forming a discrete approximation of the population vector space.

**Fig. 5:**
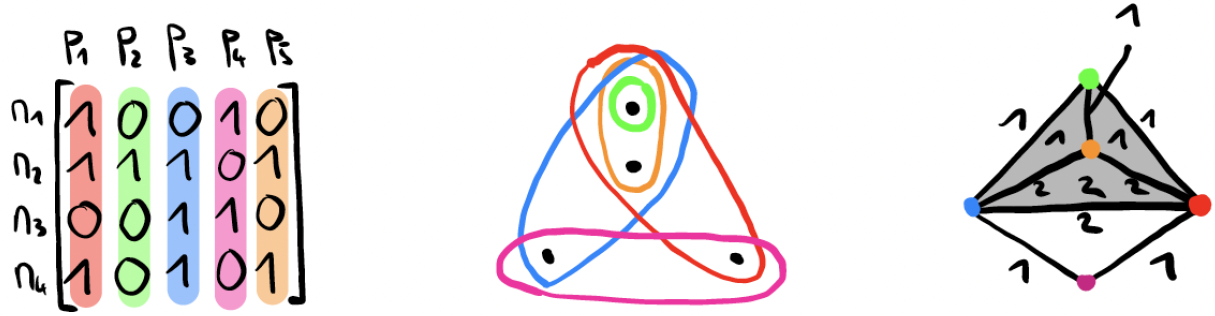
The same recording as in Fig. 4 is here color-coded according to the population vectors. Grouping neurons that are active in the same population vector (middle) gives rise to the population complex (right). The weights on the simplices indicate the number of shared neurons (equivalent to the unscaled cosine similarity).

By comparing the correlation complex (‘studying rows’, Fig 4) with the population vector complex (‘studying columns’, Fig. 5), we can then compare the two approaches of analysing a data matrix, as alluded to in the introduction. In the running example, the population vector complex has five vertices and one 3-simplex. By contrast, there are only four vertices and two 2-simplices in the correlation complex. Moreover, there is no obvious relation between the weights in the respective complexes. On the other hand, both complexes have one component (i.e., every pair of points has a path between them). Furthermore, both complexes form a loop. This might seem a lucky coincidence in this example, but, surprisingly, is not. In fact, *for every binary data matrix X, the population vector complex and the correlation complex are homotopy equivalent*.

Recall that homotopy equivalence between two spaces means we can continuously deform one space into the other without gluing or ripping. This implies that both complexes have the same number of connected components and the same number of *n*-D holes, that is, the same homology. Formally, this is due to a result in topology known as *Dowker duality*. It can be motivated intuitively [29], by first noting that the points grouped in the correlation complex are the vertices of the population vector complex (Fig. 4 and 5). That is, the population complex fits into the (good) cover given by the grouping of population vectors by neurons. From Remark 2.1 we know the combinatorics of the overlaps in this cover faithfully recovers the underlying space up to homotopy equivalence. The exact same combinatorics also yield the correlation complex of Fig. 4c. Consequently, this means that the complexes must be homotopy equivalent.

#### Remark 3.1.

If we are presented with a filtration of matrices {*X*_0_ ↪ *X*_1_ ↪…} (e.g. from applying different thresholds to a non-binary matrix of continuous firing rates), a correlation complex *CX*_*i*_ and a row complex *PX*_*i*_ may be constructed for each level of the filtration. Dowker duality holds at each level of the filtration, i.e. *CX*_*i*_ ≅ *PX*_*i*_. Furthermore, the homotopy equivalences may be chosen as to obtain a commutative diagram

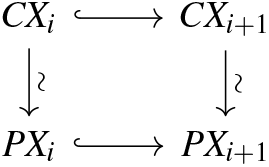

This is known as *functorial Dowker duality* [29] and implies that after applying homology, we obtain a zero-interleaving on the level of persistence diagrams. Hence, the *barcodes* associated to the filtration of correlation complexes and population vector complexes are the same. However, these filtrations do not contain information about the weights on the simplices, in which case no similar result will hold in general (see below).

## 4 Using topology to infer representations

Equipped with the constructs of Section 3, we return to the problem of inferring the covariate space of a representation directly from the data matrix *X*. First, consider the circular covariate represented in the neural activity of Fig. 6a. The recorded population vectors are sampled from a circular manifold embedded in a high-dimensional ambient space and constructing either the associated correlation complex or the population vector complex, we automatically recover the manifold up to homotopy equivalence.

**Fig. 6:**
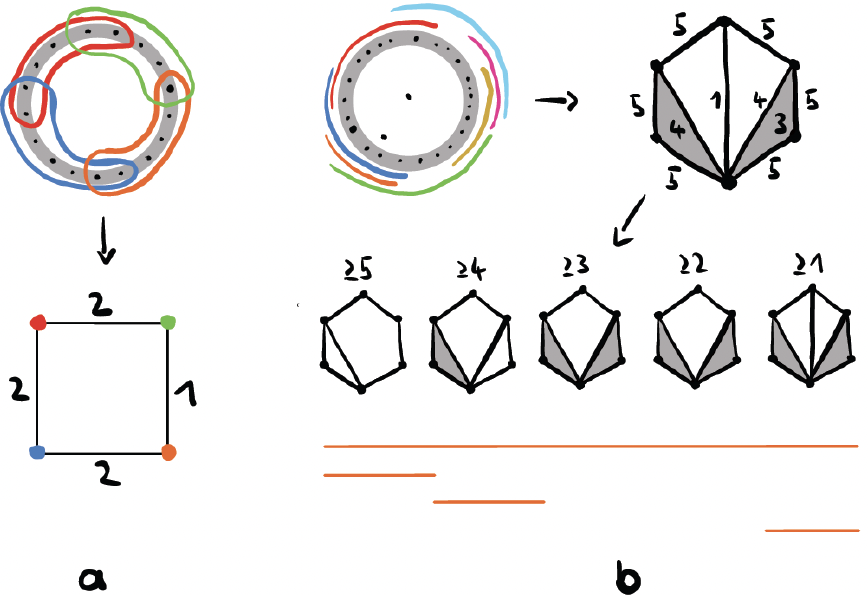
**a**. Neural activity is sampled from a neural representation of a circular covariate. **b**. Applying homology to the filtration of simplicial complexes allows determining the shape of the underlying space of the sampled points.

However, this pipeline may easily run into problems. For instance, the analysis is highly sensitive to noise – just a single spurious cofiring event could lead to a wrong result. To remedy this, we include the total weights, as introduced in Section 3, into our analysis. This is achieved by associating a filtration of (unweighted) complexes to the weighted complex (Fig. 6b). The filtration is constructed by iterating over all unique weights (from highest to lowest) and at each step adding the simplices of corresponding weight. Single topological features may then be observed at each step and are tracked over decreasing levels of the filtration. Intuitively, features persisting across many levels represent significant information, while short-lived features originate from spurious events. Formally, this analysis can be done by employing *persistent homology* [11, 30], which takes a filtration of complexes as input and outputs a *barcode* — a diagram summarizing the lifetime of topological features across the filtration (Fig. 6b).

As previously noted (Remark 3.1), and in contrast to the underlying skeleta of simplices of the correlation complex and the population vector complex, there is in general no obvious duality for the total weights. When analysing neural representations, the difference is clearly seen in two scenarios:

1. *When more than one neural ensemble is recorded*. Since two neurons of different ensembles will not have a high correlation but may only be coactive for some population vectors, we expect the correlation complex, but not the population vector complex, to depict the shape of each ensemble. Explicitly, if two ensembles represent the covariates *C*_1_ and *C*_2_, the total weights on the correlation complex will serve to infer their disjoint union *C*_1_ ⊔*C*_2_ while those on the population vector complex will rather see their product *C*_1_ *×C*_2_. An appropriate formal reconstruction result is Proposition A.4 from Appendix A.
2. *For neurons with non-convex receptive fields*. As the total weights on the population vector complex relate to the number of coactive neurons in the population vectors, the potentially ambiguous information of non-convex fields can be resolved by other neurons. Intuitively those other neurons provide a sufficently nice cover of the ambiguous non-convex fields. This information is not contained in the correlation complex. Thus, inferring the representation with the correlation complex will fail, while the population vector complex may succeed. An appropriate reconstruction result for this case is given as Proposition A.3 in Appendix A.

To illustrate the above differences, we simulated neural activity of three circular representations with each representation featuring 200 neurons (Fig. 7).

**Fig. 7:**
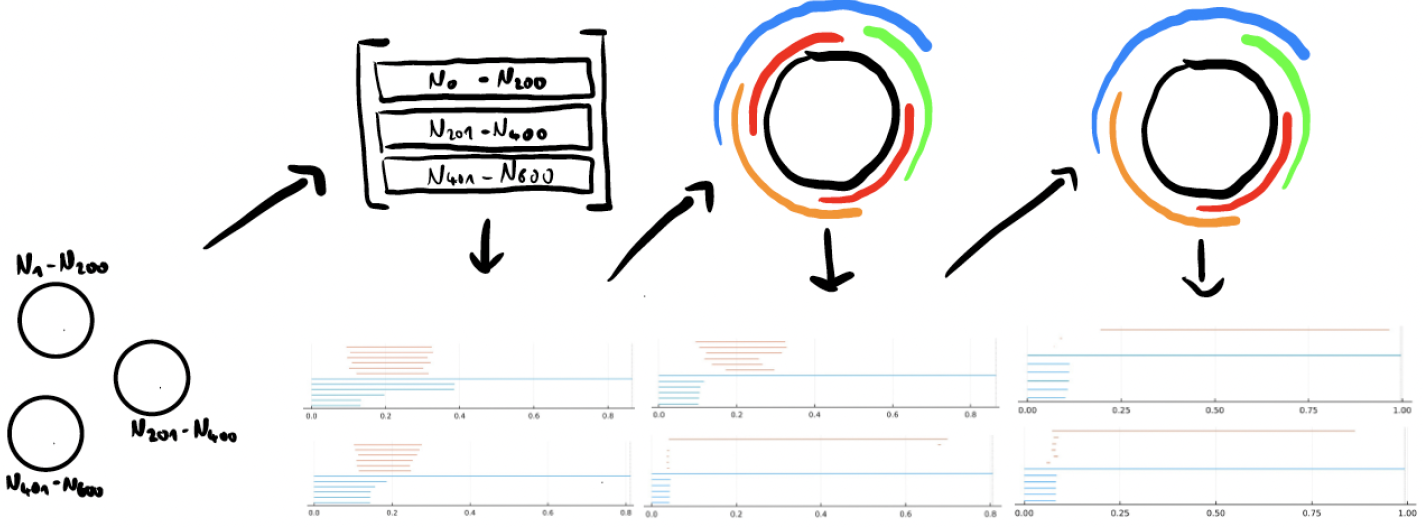
Simulation of 600 neurons with receptive fields on one of three circular covariates (far left), half of which have non-convex fields, for 1000 steps (top left). Applying homology to the correlation and population vector complex shows three long-lived 0-D features in the barcode of the former (upper bottom left) and only one in the latter (lower). This suggests clustering into three ensembles, and analysing each separately (middle). A single long-lived circular feature is visible in the barcode of the population vector complex but is obscured in that of the correlation complex due to cells with non-convex fields. Excluding these and reiterating the analysis (right), shows both barcodes feature the same topological signature.

While half of these have a single Gaussian receptive field on the respective circle, the other half has two receptive fields. Both pathologies described above are thus present, so we do not expect the correct signature of the three disjoint circles to be detected from either the correlation or the population complex. However, we can proceed by combining the information found in each:

1. In the first step the barcode of the correlation complex does not show any 1-D holes, but identifies three significant connected components.
2. Separating the data into three ensembles the population complex now yields the correct topological signature while the correlation complex is distorted by the multi-peaked fields.
3. The information of the longest-lived 1-D hole in the population complex barcode may be used to identify which neurons have multi-peaked fields [15, 31]. Excluding these, gives barcodes showing the expected significant topological features of a single circle in both the correlation complex and population vector complex.

This example shows that combining ‘studying rows’ with ‘studying columns’ may be imperative in (unsupervised) analysis of neural data, yielding results not attainable if not combined.

Converesely we may use the coincidence of topological signature, derived from the population vector complex on the one hand and the correlation complex on the other hand, as a signature for the presence of single neural modules with convex receptive fields. Showing that both approaches lead to the same inferred topology, gives greater confidence in the conclusion drawn from either. In the next section, we show how this can be of great benefit in the exploration of several (real) data sets, and how augmenting the topological analysis with cohomological decoding [15, 31] may give further insights into the internal dynamics on the inferred space.

### 4.1 Neural representations in real recordings

In this section, we further look into real data sets, showcasing the described topological inference using either the correlation complex or the population vector in recordings of spatially-tuned neurons in rodents. In Fig. 2, we already used both approaches to find a circular feature of the recorded activity in visual cortex corresponding to the orientation of the visual bars presented to the monkeys and showed that excluding the neurons with multiple peaks gave a clearer correlation structure. We note however, that neural data is inherently noisy and applying persistent homology to large data sets is computational costly [32], thus a similar preprocessing pipeline as in [17] was necessary. These preprocessing steps will influence the construction of the complexes, but the examples portray how the two approaches may be used for robust topological inference, leading to similar results when the data is primarily sampled from single functional ensembles:

- Head direction cells are neurons tuned to specific orientations the animal is facing [14]. In Fig. 8a, we analyze a 30-minute calcium imaging recording of *n* = 88 cells in the entorhinal cortex of a freely foraging mouse [33]. A ring topology is seen both in the barcodes of columns and rows of the firing rate activity matrix, as well as in the dimension-reduced projection (i,ii). Although a circular feature is detected in the correlation structure of all cells, we see that some have multiple fields on the decoded circle of the population vector complex. Excluding these in the correlation complex analyses reveals a clearer circular signature in the barcode. The represented covariate is revealed when decoding the internal dynamics and comparing it to the recorded head direction (iii, iv).
- Place cells have single firing fields in the room explored, with the combined fields of a population covering the whole environment [4]. On a 1-D, repeating virtual reality track, the cells fire at single locations in this circular environment. The environment is thus represented in the population activity, as clearly seen in the topological analysis of both rows and columns of a calcium activity matrix of *n* = 399 cells recorded in the hippocampus of a head-fixed mouse for 40 minutes [34] (Fig. 8c).
- Grid cells have multiple firing fields revealing a distinct, hexagonal pattern in 2-D space [2]. This firing pattern may be understood as each cell having a single receptive field on a toroidal state space, i.e., the activity profile is repeated in two periodic direction in correspondence with the movement of the animal [17]. Here, we analyze *n* = 152 cells in a 40-minute Neuropixel recording of a single entorhinal grid cell ensemble (also called a grid cell *module*, defined by the spacing and orientation of the firing pattern [24]) in a freely foraging rat. The toroidal structure is shown to be detectable both in the population vector and correlation complex, and its decoded coordinates correspond to the axes of the spatial pattern (Fig. 8d).

**Fig. 8:**
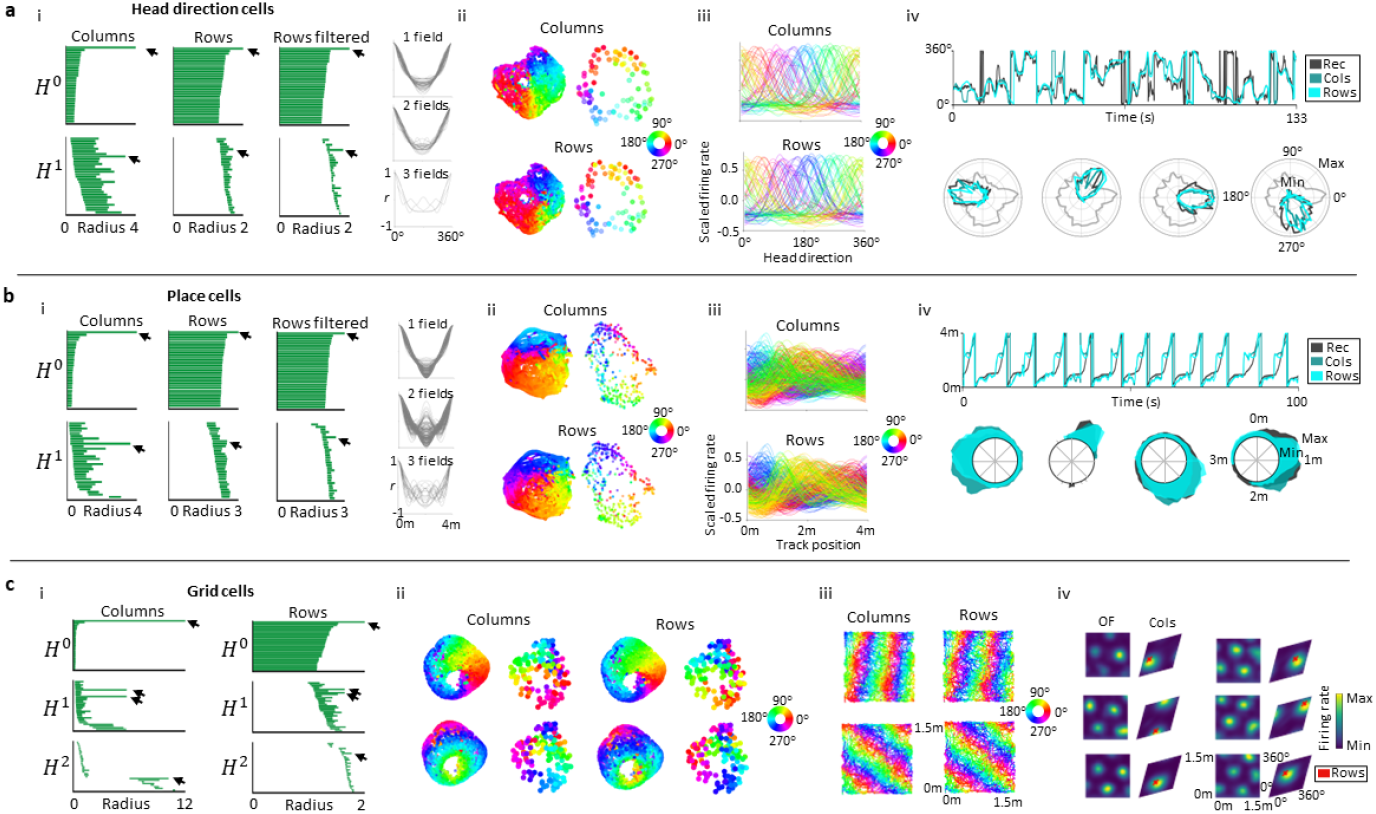
Both the population vector complex and the correlation complex reveal the topological structure of the neural representation in experimentally recorded neurons. **a**. Mouse entorhinal recordings [33] show a head direction representation. i) Left, barcodes obtained from: the population vector complex (‘Columns’); the correlation complex of all neurons (‘Rows’); the correlation complex of neurons with single tuning fields (right) on the decoded columnar circular representation (‘Rows filtered’). Arrows indicate signature topology. ii) UMAP-projection of all population vectors (left) and activity traces (right), colored according to the decoded circular coordinates from the longest-lived circular feature in either the population vector complex (top) or correlation complex (bottom). iii) Tuning curves for all neurons colored as in ii). iv) Tuning of four example neurons (bottom) and extracted time interval (top) of recorded head direction and decoded representation. **b**. Hippocampal place cells in head-fixed mice [34] uncovers a circular representation encoding the positions on the (repeated) virtual linear track. Labels and coloring as in **a. c**. Recorded activity of rat entorhinal grid cells [17] for one grid cell module shows toroidal topology in both the population vector and correlation complex. i) Barcodes as in **a-b**.i. ii) UMAP-projections as in **a-c**.ii, colored by the two periodic features of the torus (top vs. bottom), decoded from the longest-lived 1-D holes in the respective barcodes. iii) Coloring each position the rat visited by the corresponding internal representation. iv) Rate maps of six example neurons, showing tuning to spatial location in open field arena (left) and to the inferred torus (visualized as a rhombus with periodic boundaries, right). The red crosses mark the toroidal coordinates decoded from the correlation structure.

## 5 Discussion

As technological breakthroughs enable large-scale recordings of intermixed neural populations across brain regions [35], there is a need for tools to study heterogeneous neural data to understand the diverse signals. We have shown that one may study both the state space and the correlation structure through the means of topology. While these arise from the same matrix, the topological analysis of either can lead to different results – while the population vector complex exploit the strength in numbers of coactive neurons to reduce the influence of noisy or multiplexed cells, the correlation complex benefits from the persistence of coactivity in neural ensembles, undermining coincidental coactivity. We can build on both these advantages, iteratively using either approach to exhaust the information contained in discrepant results until the results coincide, at which point we obtain arobust assessment of the neural representation. Similar problems exist in gene expression analysis [36] and machine learning [37, 38], and we believe the proposed framework can be of interest beyond neuroscientific purposes. It is thus of great interest to further develop methods that integrate information obtained from ‘studying columns’ and ‘studying rows’ of a data matrix. There are many avenues to pursue this theoretically. In [39] we explain how to to leverage both directions simulataneously by constructing a bifiltration from that integrates total weights of the row and column complex in one coherent construct.

The approach of the present paper is augmented by topological decoding of the data, allowing the state space dynamics to be read out. It would be interesting to extend our framework byincorporating geometrical or statistical tools and assumptions. This is motivated both by [26] where the assumption of the precise shape of each place cell’s receptive field provides a *geometric* description of the spatial environment and [12], where the latent space is optimized both over the dynamics of the latent process as well over single cell responses to the latent space.

## Methods

### Pipeline of topological inference of real data sets

A similar setup for preprocessing the neural data as in [17] was used. This pipeline consists of combining the fuzzy topological representation constructed in the first step of UMAP [5] with persistent cohomology [11, 30]. In each case, this corresponded to computing a geodesic distance in the embedded, referred here to as the ‘fuzzy’ distance, motivated by metric realization of the fuzzy simplicial set described in [40]. The resulting distances are used in the construction of the Rips filtration in Ripser (implemented in Ripser [41, 42]), which computes the persistent cohomology, choosing 𝕫_47_-coefficients. However, certain technicalities were different in each data set in constructing the distances, as described below:

- **Head direction cells**. Calcium events (as described in [33]) sampled at 7.5Hz of mouse 97045, day 20210306, accessed in [43], were used in revealing an internal head direction representation. *Population vector complex:* The calcium events were convolved with a Gaussian filter of width *σ* = 133ms and square rooted. Time frames with speed less than *s*_0_ = 5cm/*s* or with no activity were excluded. The remaining *z*-scored population vectors were projected to their *d* = 4 first principal components and divided by the square root of the respective eigenvalues (*PCA whitening* [44]). Next, the size of the point cloud was reduced. Using the ‘fuzzy downsampling’ method introduced in [17] (setting *k* = 1000 and using Euclidean metric), left *m* = 1000 points. Finally, the negative logarithm of the ‘neighbourhood strengths’ (using cosine metric and *k* = 300, defined in the same paper), gave the fuzzy distance matrix. *Correlation complex:* A similar pipeline was used as for the population vector complex. The activity was first convolved with a Gaussian kernel of *σ* = 666ms, square rooted and time points of low speed or zero activity were excluded. The correlation distance (1 − *r*, where *r* is the Pearson correlation coefficient) between the neurons were calculated and subsequently transformed to the fuzzy distance using *k* = 65.
- **Place cells**. The deconvolved and smoothed calcium activity (30Hz sampling) found in of mouse ny226 day 20210929 was used to detect a circular representation of place cell coding. Only trials listed in ‘valid trial N’ were used. *Population vector complex:* The activity was smoothed using *σ* = 100ms and speed-filtered (*s*_0_ = 5cm/*s*). PCA-whitening was applied, projecting the data to 4 dimensions. Due to the size of the data set, a two-step downsampling procedure was performed. First, a radial downsampling scheme was used (as described in [15] with *ε* = 0.3) before the fuzzy downsampling (as previously described) reduced the point cloud to 1200 points. The fuzzy distance was computed based on the cosine metric and *k* = 900. *Correlation complex:* The correlations between the activity of all neurons (convolved as above, with *σ* = 167ms, and similarly speed-filtered), were used in computing the fuzzy distance (using *k* = 399, equal to number of neurons), to obtain the barcode of the corresponding Rips filtration.
- **V1 cells**. The spike times of the electrophysiological recordings given in ‘array 3’ of the dataset described in [18], excluding the time intervals with no visual stimuli, were used to inferring the topology of the internal representation of the orientation of drifting gratings. The spike trains were converted to firing rates by replacing each spike with a delta function (1 at time of spike and 0 otherwise) and convolving with a *σ* = 100ms Gaussian kernel. *Population vector complex:* The same pipeline as previously detailed was used, projecting the square rooted firing rates to 3 principal components and dividing by the square root of the eigenvalues, applying radial downsampling (*ε* = 0.1 and Euclidean metric) and fuzzy downsampling (*k* = 1000 and cosine metric), and computing the fuzzy distance of the remaining 1000 vectors based on the cosine metric and *k* = 500. *Correlation complex:* The persistent cohomology of the fuzzy distances between the firing rate activity of each neuron was computed based on the correlation distance and *k* = 111 (equal to the number of neurons).
- **Grid cells** The Neuropixel recordings of grid cell module 2 of Rat R day 2 as retrieved from [46] were used to find a toroidal structure both in the population vector and correlation complexes. Firing rates based on the spike times were computed as for the V1 data, using a kernel of *σ* = 50ms. *Population vector complex:* The firing rates were projected to the 7 first whitened principal components and downsampled similarly to V1 data, with radial (*ε* = 1) and fuzzy downsampling, keeping 1200 vectors. The fuzzy distance was next computed, based on cosine metric and *k* = 900. *Correlation complex:* As before, the fuzzy distances of the rows were computed based on the correlation of firing rate activity between all neurons and with *k* = 100.

### UMAP visualizations

To visualize the topological structure of the point cloud, UMAP [5] was used, procured a 3-D embedding of the data. For the ‘columns’-approach, the firing rates (as described for each data set above) was first projected to its 6 first principal components and the cosine metric and ‘n neighbours’ = 1000 was used in UMAP with otherwise default settings. The ‘rows’-visualization was similarly computed, using the correlation matrix of the firing rates and ‘min dist’ = 0.2 in the UMAP-projection.

### Cohomological decoding

To decode the circular feature(s) captured by the barcodes to compare with the recorded covariates, we used the circular coordinatization procedure introduced in [31] and with proven applicability to neural data as shown in [15, 17]. By fixing the Rips complex constructed in the Rips filtration (computed in Ripser), at filtration scale (distance) 0.99*i* + *b*, where *l* is the lifetime of the *H*^1^-bar chosen and *b* its time of birth, the corresponding cocycle representative of this circular feature assigns values in the coefficient ring (in this case ℤ_47_) to all edges (i.e., 1-simplices) in the complex. In essence, the method then smooths these values by minimizing their variation across all edges. This allows us a map from each vertex to the circle, i.e. providing circular coordinates to each point (in our case corresponding to either a population vector or the activity of a single neuron). Hence, this gives a decoding of the circular feature represented in the neural activity.

When constructing the population vector complex, the point cloud was downsampled, and hence, not all points were assigned a coordinate from the above-described method. Thus, the coordinates obtained were extrapolated to the whole (dimension-reduced) point cloud by first computing a distribution for each principal component by weighing each coordinate by the vector elements of the corresponding time frames. For each vector in the remaining point cloud, each of the distributions were weighted in a similar manner as above and summed to give a single distribution for each point whose mass center then defined the desired coordinate.

Similarly, in finding time-varying angles based on the decoded coordinates of the correlation complex, the coordinates were weighed by the firing rate activity of the population vector at each time frame and the mass center computed to get the coordinate of highest mean activity.

Likewise, the coordinates assigned to each neuron based on the decoded population vectors were computed as the mass centers of each neuron’s firing rate tuning to the decoded coordinates.

The coordinate sets were aligned by choosing the orientation and origin with lowest mean angular difference to the recorded covariate.

### Detection of multiple receptive fields

To find the number of tuning fields per neuron in the relevant data sets, the average firing rate activity was found in even bins of 6^°^ of the decoded circular coordinates of the population vectors. The resulting distribution was convolved with a Gaussian kernel of 18°-width using periodic boundary conditions and subsequently normalized to 0–1. Horizontal lines were then defined (*x*-axis describing the decoded circular coordinate and *y* the normalized firing rate) for 60 evenly spaced values of *y*. The number of intersections between the lines and the normalized firing rate distribution, found using the python package ‘shapely.LineString’, was counted and divided by two. Rounding this number to its lowest integer gave the number of ‘bumps’ the line crossed. The maximal number of bumps detected across each line was used to determine the number of tuning fields for each neuron. This allowed inferring the topology of the correlation complex with distinct (single-bumped) tuning to the column-wise circular representation of the ensemble.

## Acknowledgments

We are grateful to Graf et al [18], Weijian Zong et al [33], Gardner et al [17], Pettit et al [34] for sharing data sets and to Žiga Virk for helpful conversations that in particular motivated the formualtion of Propositions A.3 and A.4.

## Appendix A Formal background and reconstruction results

This appendix seeks to give some formal background on the observations and constructions from the main body of the text. In particular we state and prove the two reconstruction results alluded to in Section 4.

### Dowker complexes and total weights

In [47] Dowker introduced the row and column complex of a relation *A* : *I × J* → {0, 1}. To be in tune with the main body of this paper let us think of such a relation as a binary matrix *A* with indices *I×J* where *A*_*i, j*_ = *A*(*i, j*) iff (*i, j*) ⊆ *A*. The row complex *R*(*A*) has simplices given as {*σ* ⊆ *I* | ∃ *j* ∈ *J*, ∀*i* ∈ *σ* : *A*(*i, j*) = 1} i.e by subsets of *I* collectively satisfying the relation with some elements of *J*. Similarly the column complex of *C*(*A*) has simplices {*τ* ⊆ *I* | ∃*i* ∈ *I*, ∀ *j* ∈ *τ* : *A*(*i, j*) = 1}. Dowker proved that *R*(*A*) and *C*(*A*) always have isomorphic simplicial homology groups. This was later generalized by Björner to the following statement.

#### Theorem A.1.

For every relation *A*, the realizations of the row and column complex are weakly homotopy equivalent.

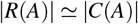

We may weight the simplices of the row and column complex by their numbers of witnesses. In [48] explains that these total weights give us filtrations of the row complex

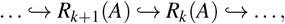

where *σ* is a simplex in *R*_*k*_(*A*) if it has at least *k* witnesses i.e #{*j* ∈ *J* | *A*_*i*, *j*_ = 1} ≥ *k*. Similarly we obtain a filtration of the column complex

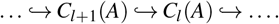

### The nerve and the Vietoris complex of a cover

Given an open cover 𝓊 = {*U*_*i*_ ⊆ *X* }_*i*∈*I*_ of a topological space *X* we can form it’s *nerve complex* given as N𝓊 with simplices {*σ* ⊆ *I* | _*i*∈*σ*_ *U*_*i*_*≠* ∅}. If all intersections of elements in the cover 𝓊 are either contractible or empty we call it a *good* cover. Then all homotopiocal information about the space *X* is concentrated in the combinatorics of which cover elements overlap. This information is encoded in the nerve complex and the following classic result, known as the *nerve lemma* comes as no surprise.

#### Theorem A.2.

Let 𝓊 be a good open cover of the topological space *X*. Then

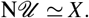

Given an open cover 𝓊 = {*U*_*i*_ ⊆ *X*}_*i*∈*I*_ we may define a relation *A*_𝓊_ ⊆ *I* ×*X* with (*i, x*) ∈ *A*_𝓊_ iff *x* ∈ *U*_*i*_. The row complex of this relation is precisely the nerve of the cover 𝓊. Also it’s column complex has a name. It’s the covers *Vietoris complex* V𝓊.

### Two reconstruction results

Let 𝓊 = {*U*_*i*_ ⊆ *X* }_*i*∈*I*_ be an open cover of *X* and *J* ⊆ *X* a sample of points such that all non-empty intersections *U*_*σ*_ = ∩_*i*∈*σ*⊆*I*_*U*_*i*_ are witnessed by some element of *J*. Define

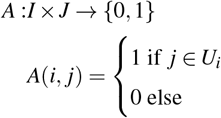

and let *R*_*•*_(*A*) and *C*_*•*_(*A*) be the filtrations of row and column complexes of this relation in terms of total weights as defined above.

#### Proposition A.3.

Let *m* ≤ *n*. If

1. 𝓊 has multiplicity at least *n* and
2. *U*_*σ*_ is either empty or contractible for every *σ* ⊆ *I* with #*σ* ≥ *m*, then for *m* ≤ *k* ≤ *n*: |*C*_*k*_| ≅ *X*.

*Proof*. Define *I*^*k*^ = {*σ* ⊆ *I* |#*σ* ≥ *k*} and then 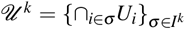. Because of assumption (1) we know that 𝓊^*k*^ is an open cover of *X* for *k* ≤ *n*. If *m* ≤ *k* then all intersections of cover elements satisfy assumption (2) and are thus contractible or empty: the cover is good. Define a new relation

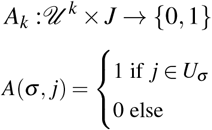

It’s row complex is the nerve N*𝓊* ^*k*^ and thus homotopy equivalent to *X*. The column complex is precisely *C*_*k*_(*A*). An application of Dowker duality (Theorem A.1) finishes the proof.

Now let 𝓊 _1_ = {Ũ_*i*_ ⊆ *X*_1_}_*i*∈*I*1_ and 𝓊 _2_ = {Ũ_*i*_ ⊆ *X*_2_}_*i*∈*I*2_ be two *good* open covers. Let *I* = *I*_1_ ⊔*I*_2_ and define 𝓊 = {*U*_*i*_ ⊆ *X*_1_ *×X*_2_}_*i*∈*I*_ with

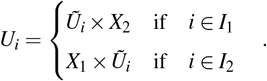

Then let *J* ⊆ *X ×Y* and define the usual relation *A* : *I ×J* → {0, 1} indicating membership of sample points to cover elements.

#### Proposition A.4.

Let *m* ≤ *n*. If

1. For all *σ* ⊆ *I*_1_ or *σ* ⊆ *I*_2_ with *U*_*σ*_ non empty we have #(*U*_*σ*_ ∩*J*) ≥ *n* and
2. for all (*i*_1_, *i*_2_) ∈ *I*_1_ *×I*_2_ we have 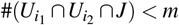, then for all *m* ≤ *k* ≤ *n*: |*R*_*k*_| ≅ *X* ⊔*Y*

*Proof*. It is fairly obvious that *R*_*k*_(*A*) decomposes into two connected components that are the nerves N𝓊 _1_ and N𝓊 _2_ respectively. Just to illustrate the similarity with the situation of Proposition **??** we give a more complicated proof.

For *j* ∈ *J* define *I*_*j*_ = {*i* ∈ *I*| *j* ∈ *U*_*i*_} and then *𝒱*_*k*_ = {*V*_*τ*_ = ∩_*j*∈*τ*_ *I*_*j*_|#*τ* ≥ *k* }. Because of assumption (2) all *V*_*τ*_ ∈ *𝒱*_*k*_ are either contained in *I*_1_ or in *I*_2_. We may think of them as points in *X* ⊔*Y*. Moreover we derive from assumption (1) that all non-empty intersections ∩_*i*∈*σ*⊆*I*1_Ũ_*i*_ and ∩_*i*∈*σ*⊆*I*2_*Ũ*_*i*_ are witnessed by some element of *𝒱*_*k*_ i.e there is *V*_*τ*_ ⊆ *σ*. Define a new relation

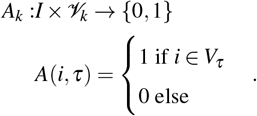

The column complex of this relation is the Vietoris complex of the cover 𝓊_1_ ∪ 𝓊_2_ and thus homotopy equivalent to *X* ⊔*Y*. The row complex of *A*_*k*_ is precisely *R*_*k*_(*A*) and again we finish the proof by an application of Dowker duality (Theorem A.1).

